# An EWS::FLI1 monoclonal antibody with multimodal utility and specificity for FLI1

**DOI:** 10.1101/2023.11.13.566886

**Authors:** Saravana Selvanathan, Jeffrey Toretsky

## Abstract

Ewing sarcoma (ES) is a rare tumor that affects children, adolescents, and young adults. This tumor has a high morbidity in all patients and for those who present with metastatic disease a high mortality. A canonical chromosomal translocation, either t(11;22)(q24;p12) or t(21;22)(q22;q12) leads to the fusion oncoproteins EWS::FLI1 or EWS::ERG in 95% of ES patients. We recognized a critical need for a stably sourced high affinity antibody that recognizes EWS::FLI1 with maximal specificity. We created and validated a hybridoma that produces a mouse derived monoclonal antibody that recognizes EWS::FLI1 in multiple molecular biology applications.

## Introduction

Ewing sarcoma (ES) is a rare tumor that affects children, adolescents, and young adults. This tumor has a high morbidity in all patients and for those who present with metastatic disease a high mortality. A canonical chromosomal translocation, either t(11;22)(q24;p12) or t(21;22)(q22;q12) leads to the fusion oncoproteins EWS::FLI1 or EWS::ERG in 95% of ES patients. These fusion proteins are critical initiators and maintenance proteins in the patient tumors. They uniquely occur in the tumor and not the non-cancer tissue so their specific presence and vital nature in ES make them an ideal therapeutic target. A wide range of biologic experiments requires having an antibody that can recognize these proteins in a variety of assays including immunoblotting, immunohistochemistry, immunoprecipitation, and chromatin immunoprecipitation. Detection of native proteins adds significant validity to conclusions about ES biology.

Since the discovery of these fusion oncoproteins, a reproducible source of antibodies has been desired. Early polyclonal antibodies were often useful but limited by the production in a given rabbit or other animal. While monoclonal antibodies would be ideal if derived from a renewable hybridoma, these have not been commercially available. We undertook a project to create a human monoclonal antibody that would be useful in a variety of assays synthesized in a renewable hybridoma.

### Materials and Methods Hybridoma cell culture

Hybridoma cells were grown in DMEM with 15% FBS at 37°C in 5% CO2. Healthy expansion cells were split at 60% confluence. Hybridoma cells were removed from the plate by gentle pipetting and do not need trypsin digestion. Change media 2-3 times per week. When cell density reaches 1×10^6^/ml, supernatants were collected after 12 hrs of incubation.

### Cell lines and reagents

ES cell lines TC32 and A4573, were grown in RPMI with 10% FBS and 1% HEPES. All cell lines were grown at 37°C in 5% CO_2_ and ES cells were passaged every 2–4 days. Cell line integrity was confirmed by fingerprinting. Cell lines were tested for mycoplasma *in domo* at regular intervals.

### Nuclear co-immunoprecipitation

Co-immunoprecipitation (CoIP) was performed to validate EWS::FLI1 complex protein-protein interaction. Active motif universal magnetic Co-IP kit (Active Motif) was used to make nuclear extract from TC32 and A4573 cells according to the manufacturer’s instructions. The protein G magnetic beads were used for Co-IP and the IP was performed on 300 μg samples using 2 μg of custom FLI1 monoclonal antibody and mouse monoclonal IgG (as a negative control). Western blots were performed individually using the following antibodies: RHA (Everest Biotech #EB09297), PRPF8 (abcam # ab79237), SFPQ (abcam # ab99357), hnRNPK (abcam # ab18195), and SRSF3 (abcam # ab73891). Detection was carried out using Millipore Immobilon Western Chemiluminescent HRP Substrate per the manufacturer’s instructions (Millipore Corp.) using a Li-COR Odyssey FC Imaging System.

### Western blot

Samples were subjected to SDS-PAGE and then transferred to a Immobilon-P membrane (Millipore, Billerica, MA, USA). Non-specific binding sites were blocked upon incubation in 5% nonfat dry milk diluted in 1X TTBS (20 mM Tris-HCl, pH 7.5, 150 mM NaCl, 0.05% Tween 20) for 1 h at RT. The membranes were incubated with primary antibodies diluted in 5% BSA in 1X TTBS either at RT for 2 h or at 4°C overnight according to manufacturer’s recommendations. Dilution for primary antibodies FLI1, PRPF8, RHA, SFPQ, hnRNPK and SRSF3 were at 1:1000 and anti-actin-HRP at 1:5000 (Santa Cruz Biotechnology, Santa Cruz, CA, USA; #sc-1615). After rinsing three times with 1X TTBS, the membranes were incubated for 1 hr at RT in HRP conjugated anti-rabbit (GE Healthcare Bio Sciences, Pittsburgh, PA, USA; #NA934V) or anti-mouse (GE Healthcare Bio Sciences, Pittsburgh, PA, USA; #NA931V) secondary antibody diluted 1:5000 in 1X TTBS or anti-goat secondary antibody (Santa Cruz Biotechnology, USA; # sc-2354) diluted 1:3000 in 5% nonfat dry milk in 1X TTBS. The blots were then washed three times in 1X TTBS and then developed using Millipore Immobilon Western chemiluminescent HRP substrate according to the manufacturer’s instructions (Millipore Corporation, Billerica, MA, USA).

### Dot blot

Similar to the western blot technique but differing in that protein samples are not separated electrophoretically but are spotted through circular templates directly onto the membrane. Using a narrow-mouth pipette tip, spot 2 μl of samples onto the nitrocellulose membrane at the center of the grid. Minimize the area that the solution penetrates (usually 3-4 mm diam.) by applying it slowly and let the membrane dry. Non-specific binding sites were blocked upon incubation in 5% nonfat dry milk diluted in 1X TTBS (20 mM Tris-HCl, pH 7.5, 150 mM NaCl, 0.05% Tween 20) for 1 h at RT. The membranes were incubated with primary antibodies diluted 1:1000 in 5% BSA in 1X TTBS at RT for 2 h. After rinsing three times with 1X TTBS, the membranes were incubated for 1 h at RT in HRP conjugated anti mouse (GE Healthcare Bio Sciences, Pittsburgh, PA, USA; #NA931V) secondary antibody diluted 1:5000 in 1X TTBS. The blots were then washed three times in 1X TTBS and then developed using Millipore Immobilon Western chemiluminescent HRP substrate according to the manufacturer’s instructions (Millipore Corporation, Billerica, MA, USA). Chemiluminescence was detected using a Li-COR Odyssey FC imaging system.

### Surface plasmon resonance (SPR)

The SPR experiments were performed using a Biacore T200 SPR instrument (GE Healthcare) with CM5 chip at 25 C. Anti-FLI1 Antibody (∼150000Da, 1.0 mg/ml stock) was used as the ligand to capture onto CM5 sensor surface. EWS-FLI1 (∼68000 Da, 1.7 mg/ml stock) was used as analytes to flow over the ligand immobilized surface in the absence and presence of Peptide-1 (∼1440.6 Da, 10 mM stock) captured on the antibody surface. Flow Cell (FC) 3 was used as reference for FC4. Anti-FLI1 Antibody was diluted (1:100, 0.01 mg/mL) in 10 mM sodium acetate buffer at pH 5.5 and immobilized onto FC4 to a level of ∼3500 RU, using standard amine coupling chemistry. HBS-P (10 mM Hepes pH 7.4, 150 mM NaCl, 0.05% surfactant P20) was used as the immobilization running buffer. Based on the capture response values, theoretical R_max_ values were calculated.

The R_max_ values assume 1:1 binding for all analytes. Overnight kinetics was performed for all analytes binding to the ligand in the presence of HBS-P+1%DMSO+0.1%Glycerol. The flow rate of all analyte solutions was maintained at 50 μL/min. The contact and dissociation times used were 60s and 300s, respectively. EWS-FLI1, FLI1, and ERG analyte concentrations were from 100 nM to 6.25 nM (four-fold dilutions). These concentrations were injected in the presence and absence of Peptide-1 and Peptide-4 captured on the antibody surface. 1000 nM of each peptide was injected to saturate the antibody surface whenever the capture of peptides required. Two 20 s pulses of 50 mM NaOH were injected for surface regeneration. All analytes were injected in duplicate. Sensorgram from the overnight kinetics were plotted for further evaluation.

### Nikon SoRa Spinning Disk Microscope

Cells were seeded at 100,000 cells per well in 12 well plates on sterilized, collagenized coverslips. After 24 hours, the coverslips were fixed with 10% formalin for 5 minutes and washed with Phosphate Buffer Saline (PBS). The coverslips were then treated with 0.5% Triton X-100 for 15 minutes to permeabilize the nucleus and washed 3 times with PBS. 5% Bovine Serum Albumin (BSA) was added for a 1-hour incubation to eliminate non-specific binding, and a signal enhancer was added for 30 minutes to reduce charge-linked fluorescent background. Overnight incubation with primary antibody diluted in 0.5% BSA was followed by 3 PBS rinses, and a fluorophore-conjugated secondary antibody incubation for 1 hour in darkness. The coverslips were then rinsed with PBS and treated with 300 nM DAPI solution for 5 minutes, followed by mounting with Diamond Prolong Antifade Mountant onto glass slides. After curing for 48 hours, the slides were imaged at 60X magnification with immersion oil on a Nikon SoRa Spinning Disk Microscope under the following conditions: DAPI nuclear counterstain (λex 350, λem 470) Mouse monoclonal EWS::FLI1-Goat anti-mouse AlexaFluor 488 (λex 490, λem 525) Rabbit monoclonal hnRNPK-Goat anti-rabbit AlexaFluor 594 (λex 590, λem 617). The images were processed with denoise and deconvolution NIS software. Colocalization analysis was performed using BIOP JACoP (Bioimaging and Optics Platform/ Just Another Colocalization Plugin) through ImageJ.

## Results

We selected a series of contiguous amino acids and created 12-mer peptides for immunization into mice. Hybridomas were created and ascites was tested against lysate from Ewing sarcoma (ES) cell lines A4573 and TC32. Recombinant full-length EWS::FLI1 was used a a control and HEK293 as a negative control (Figure 1A). Dot blot titrations of ES lysate showed binding with all antibodies, however 1.1 and 1.2 had the highest titers (Figure 1B). Further screening occurred with an immunoprecipitation of RNA Helicase A (RHA) showing that peptide 1.1 demonstrated promising activity (Figure 1C).

**Figure 1:**
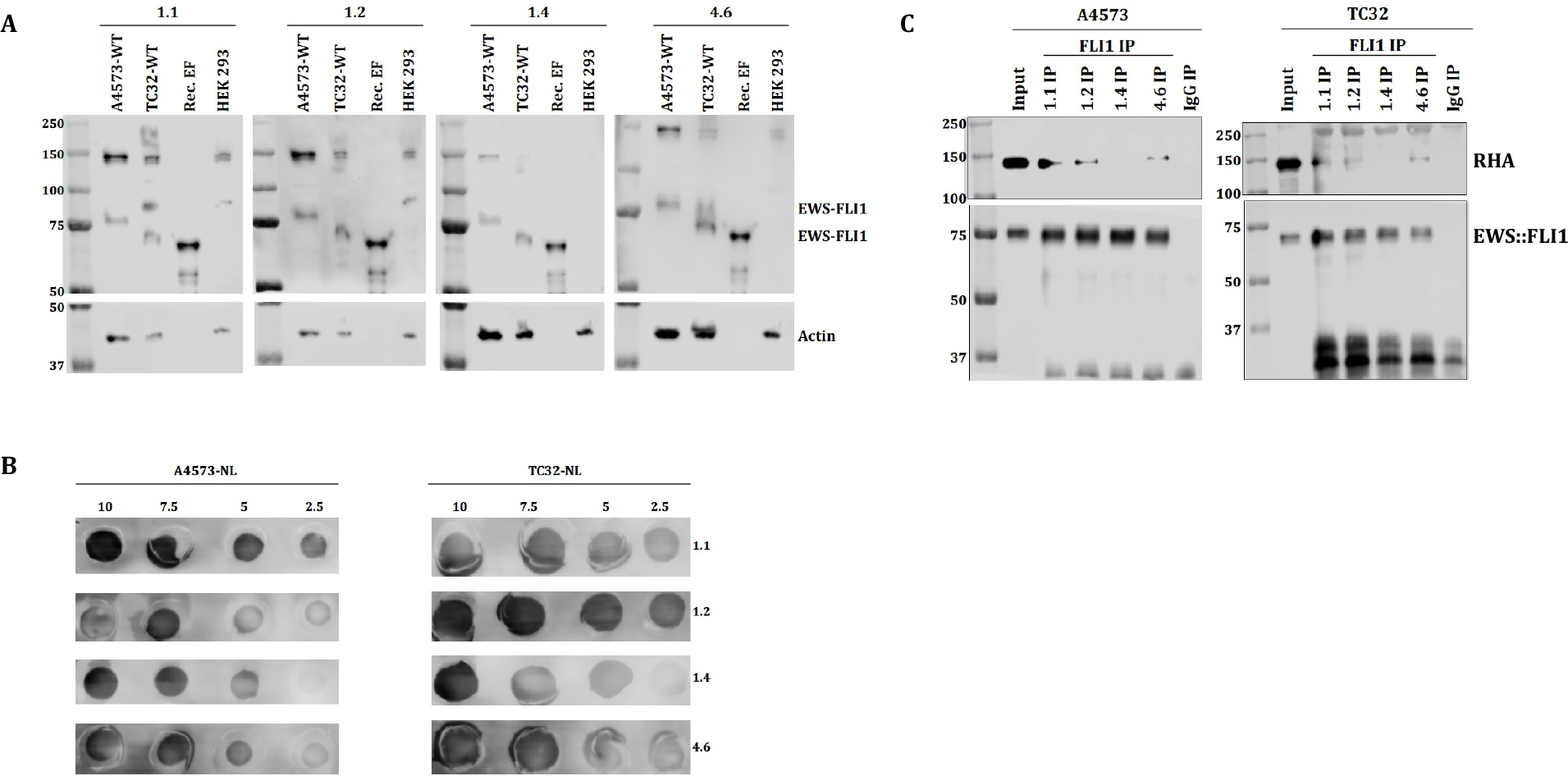
Selected hybridoma titration and antibody binding to EWS::FLI1. (**A)** Western blot analysis showing selected custom FLI1 mAb detection of EWS::FLI1 fusion protein in the ES cell line TC32 (type 1) and A4573 (type 3), recombinant EWS::FLI1fusion protein used as a positive control, and HEK293 used as a negative control. Actin used as a loading control. (**B)** Dot blot analysis showing different concentration of ES nuclear lysate detection with selected FLI1 mAb hybridoma clones. (**C)** Nuclear co-immunoprecipitation (co-IP) of EWS::FLI1 and RHA using selected custom FLI1 mAb. 10% of total nuclear lysate was used as input, selected FLI1 antibody precipitates show the presence of proteins as a part of the complex. For the negative control, mouse monoclonal IgG antibody was used to show specificity.

Immunopurification allowed for concentration of the antibody and performed for the peptide 1.1 antibody prior to further testing. Antibody binding was confirmed with surface plasmon resonance, initially showing that the immunization peptide 1.1 had a KD for the antibody of 20 pM (Figure 2A). Full length recombinant EWS::FLI1 bound with a KD of 300 nM (Figure 2B). Peptide competition showed that the full length EWS::FLI1 was displaced by the peptide in binding to the antibody (Figure 2C).

**Figure 2.**
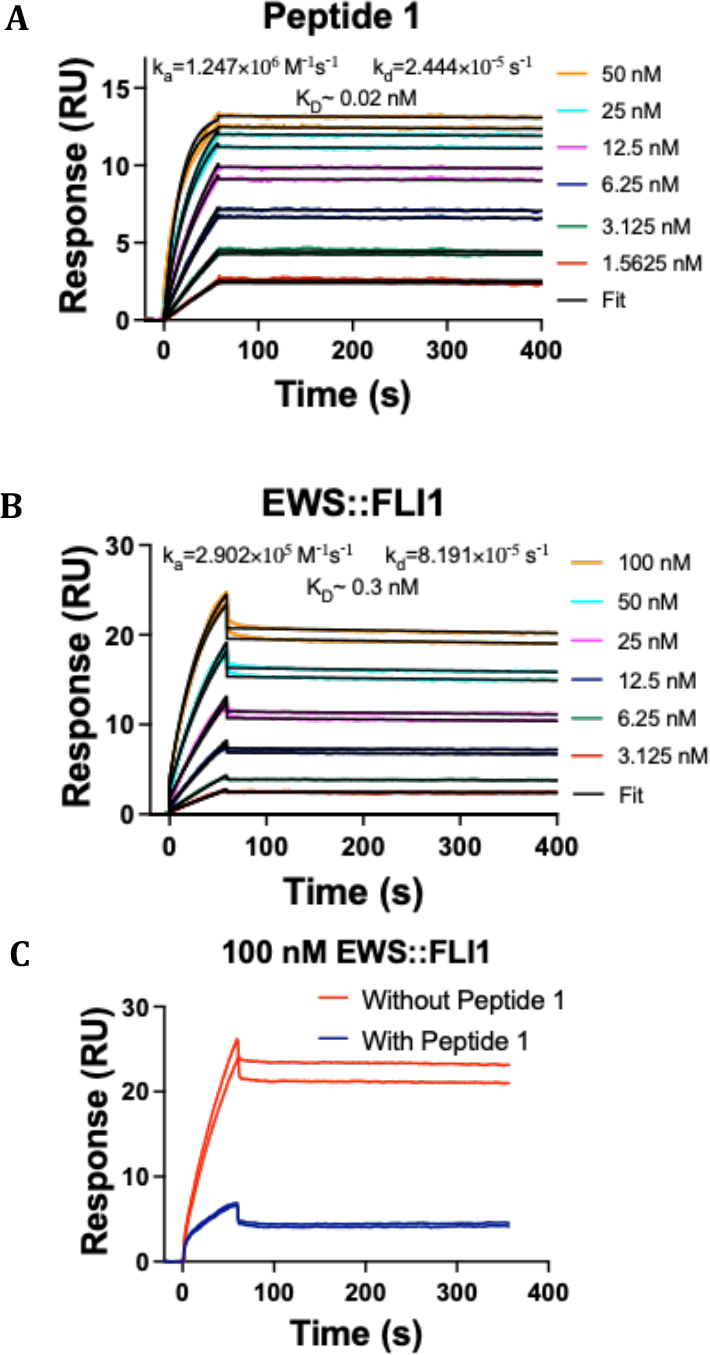
Surface plasmon resonance confirms antibody specificity and effective peptide competition. (**A)** SPR sensorgram showing binding of Peptide 1 to immobilized custom FLI1 mAb. Colored lines are experimental data and black lines are 1:1 kinetics fit. (B)SPR sensorgram showing binding of EWS::FLI1 to immobilized custom FLI1 mAb. Colored lines are experimental data and black lines are 1:1 kinetics fit. (C)SPR sensorgram showing binding of 100 nM EWS::FLI1 to over immobilized custom FLI1 mAb in the presence and absence of Peptide 1 injected over the custom FLI1 mAb immobilized surface. Data shows a 1:1 kinetics fit.

Immunopurified Antibodies 1.1 and 1.4 were tested in immunoblots with two ES cell lines and compared with blots incubated in the presence of the immunizing peptide. Good signals are seen for the 75 kDa EWS::FLI1 type 3 protein in the A4573 cells lysates and the 65 kDa EWS::FLI1 type 1 protein in TC32 (Figure 3A). Both bands are competed from the peptide and the HEK cells do not show any product as a negative control. Detecting EWS::FLI1 with immunohistochemistry and confocal microscopy shows strong nuclear localization (DAPI stain) with a diminished signal as the peptide is titrated into the incubating mixture (Figure 3B).

**Figure 3:**
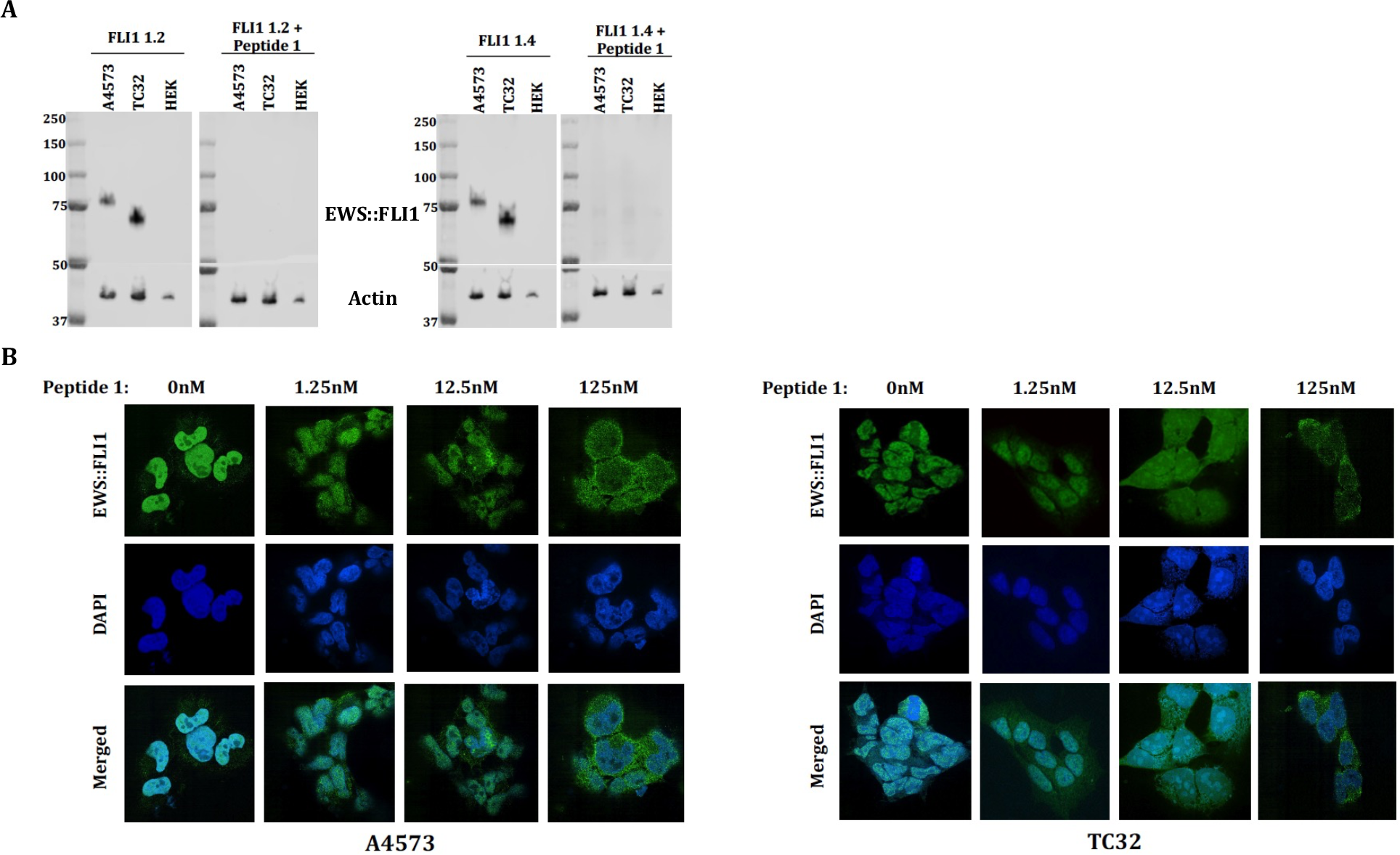
Validation of EWS::FLI1 custom mAb demonstrates specific fusion protein binding in ES cells with strong immunizing peptide competition. (**A)** Western blot analysis showing EWS::FLI1 fusion protein detection in the ES cell line TC32 (type 1) and A4573 (type 3), HEK293 used as a negative control and the addition of the peptide 1 containing the same epitope as EWS::FLI1. Actin was used as a loading control. (**B)** Coverslips from A4573 or TC32 were incubated with Antibody 1 with/without Peptide 1. The nuclear signal is extinguished by 125 nM and 12.5 nM peptide for cell lines A4573 and TC32 respectively. Images taken at 60x on Nikon SoRa Spinning Disk Microscope.

Immunoprecipitation of EWS::FLI1 with protein partners is a critical method for understanding protein function. We show that Antibody 1.1 successfully co-immunoprecipitated EWS::FLI1 partners PRPF8, RHA, SFPQ, hnRNPK, and SRSF3 in both A4573 and TC32 (Figure 4A). Using IHC, Antibody 1 is able to effectively identify EWS::FLI1 and show that it colocalizes with hnRNPK (Figure 4B).

**Figure 4:**
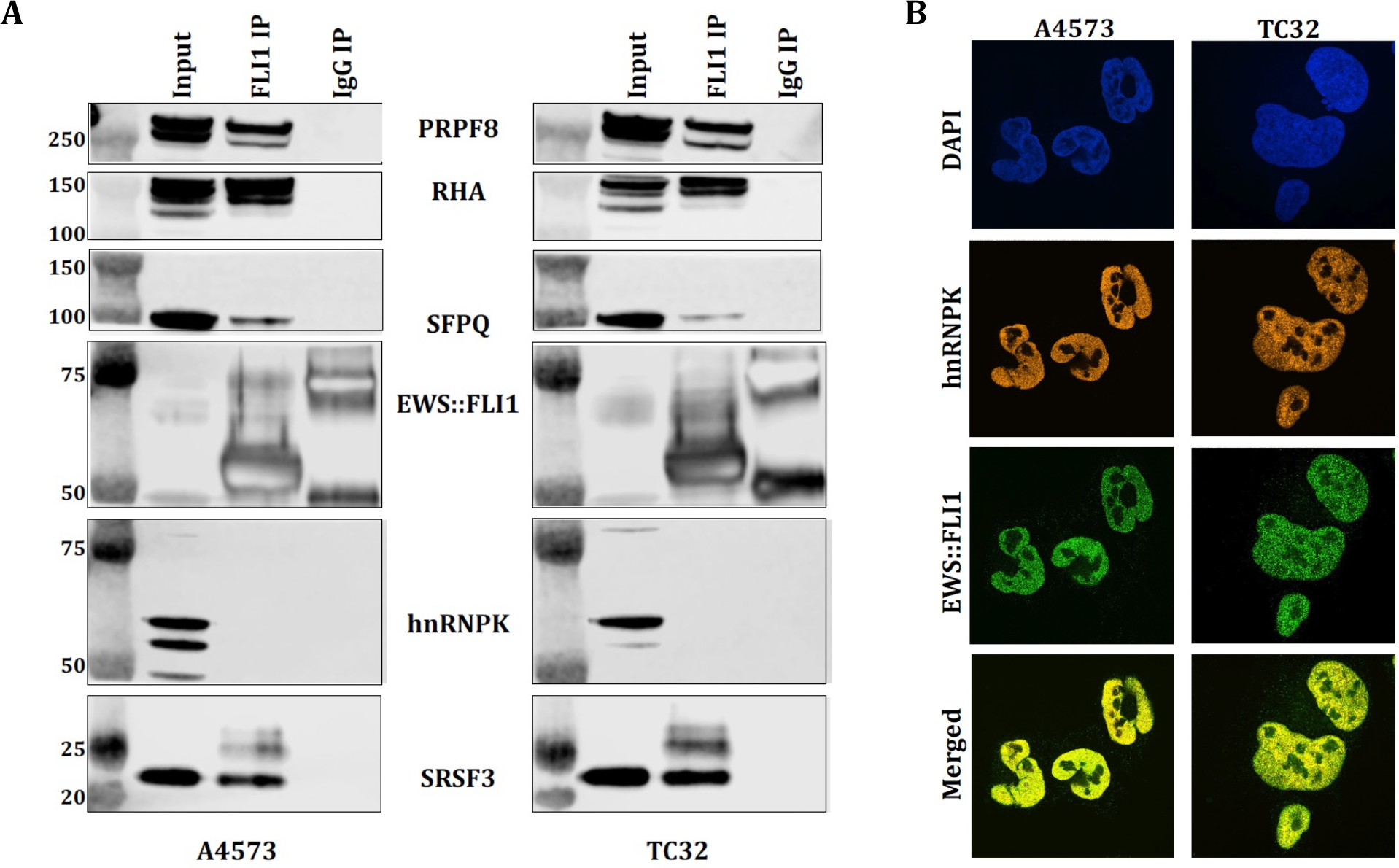
Purified antibody shows effective uses for EWS::FLI1 co-immunoprecipitation and co-localization. **(A)** Antibody 1 immunoprecipitation of EWS::FLI1 with a series immunoblots to identify co-immunoprecipitated proteins. Protein markers (Lane 1), 10% of total nuclear lysate was used as input (lane 2), custom FLI1 mAb precipitates show the presence of proteins as a part of the complex (lane 3). For the negative control, mouse monoclonal IgG antibody was used to show specificity (lane 4). (**B)** Immunohistochemistry of ES coverslips stained with Antibody 1 to identify EWS::FLI1 in ES cell line A4573 and TC32 and stained with hnRNPK antibody. Both A4573 and TC32 split into channels DAPI (blue), hnRNPK (orange), and EWS::FLI1 (green) imaged at 60x on a Nikon SoRa Spinning Disk Microscope. Merged channels (yellow) show the overlap between hnRNPK and EWS::FLI1, depicting colocalization between the two proteins.

## Discussion

We show a successful production of a new antibody that has remarkable specificity and affinity for EWS::FLI1. The multiple applications of this antibody support its broader utility. Further studies of paraffin tissue and additional microscopy are pending at this time.

## Notes

### Competing Interest Statement

The authors have declared no competing interest.

